# Chondroitin sulfate glycan sulfation patterns influence histochemical labeling of perineuronal nets: a comparative study of interregional distribution in human and mouse brain

**DOI:** 10.1101/2024.02.09.579711

**Authors:** Claudia Belliveau, Stéphanie Théberge, Stefanie Netto, Reza Rahimian, Gohar Fakhfouri, Clémentine Hosdey, Maria Antonietta Davoli, Aarun Hendrickson, Kathryn Hao, Bruno Giros, Gustavo Turecki, Kimberly M. Alonge, Naguib Mechawar

## Abstract

Perineuronal nets (PNNs) are a condensed subtype of extracellular matrix that form a net-like coverings around certain neurons in the brain. PNNs are primarily composed of chondroitin sulfate (CS) proteoglycans from the lectican family that consist of CS-glycosaminoglycan (CS-GAG) side chains attached to a core protein. CS disaccharides can exist in various isoforms with different sulfation patterns. Literature suggests that CS disaccharide sulfation patterns can influence the function of PNNs as well as their labeling. This study was conducted to characterize such interregional CS disaccharide sulfation pattern differences in adult human (N = 81) and mouse (N = 19) brains. Liquid chromatography tandem mass spectrometry was used to quantify five different CS disaccharide sulfation patterns, which were then compared to immunolabeling of PNNs using *Wisteria Floribunda Lectin* (WFL) to identify CS-GAGs and anti-aggrecan to identify CS proteoglycans. In healthy brains, significant regional and species-specific differences in CS disaccharide sulfation and single versus double-labeling pattern were identified. A secondary analysis to investigate how early-life stress (ELS) impacts these PNN features discovered that although ELS increases WFL+ PNN density, the CS-GAG sulfation code and single versus double PNN-labeling distributions remained unaffected in both species. These results underscore PNN complexity in traditional research, emphasizing the need to consider their heterogeneity in future experiments.

## Introduction

Perineuronal nets (PNNs), first described by Camillo Golgi in 1893, are a condensed and specialized form of extracellular matrix (ECM) that surround the soma and proximal dendrites of various neurons in the central nervous system (Novak, U. and Kaye, A.H. 2000). PNNs form preferentially around parvalbumin (PV) expressing inhibitory interneurons and, to a lesser extent, excitatory neurons (Tanti, A., Belliveau, C., et al. 2022). Comprising mainly chondroitin sulfate (CS) proteoglycans (CSPGs) from the lectican family (aggrecan, brevican, neurocan and versican), PNNs are synthesized by both the surrounding glia and neurons themselves exhibiting a highly organized mesh-like structure (**Figure 1A**) (Carulli, D., Rhodes, K.E., et al. 2006, Celio, M.R., Spreafico, R., et al. 1998, Schwartz, N.B. and Domowicz, M.S. 2018, Tanti, A., Belliveau, C., et al. 2022). The CSPGs are attached at their N-terminus through link proteins to a hyaluronan backbone that is secreted by hyaluronan synthase (HAS) located on the cell surface (Kwok, J.C., Dick, G., et al. 2011) (**Figure 1B**). The CSPGs then bind to one another through Tenascin-R at their C-terminus, thus forming their characteristic PNN ‘net-like’ structures (Lundell, A., Olin, A.I., et al. 2004).

**Fig. 1:**
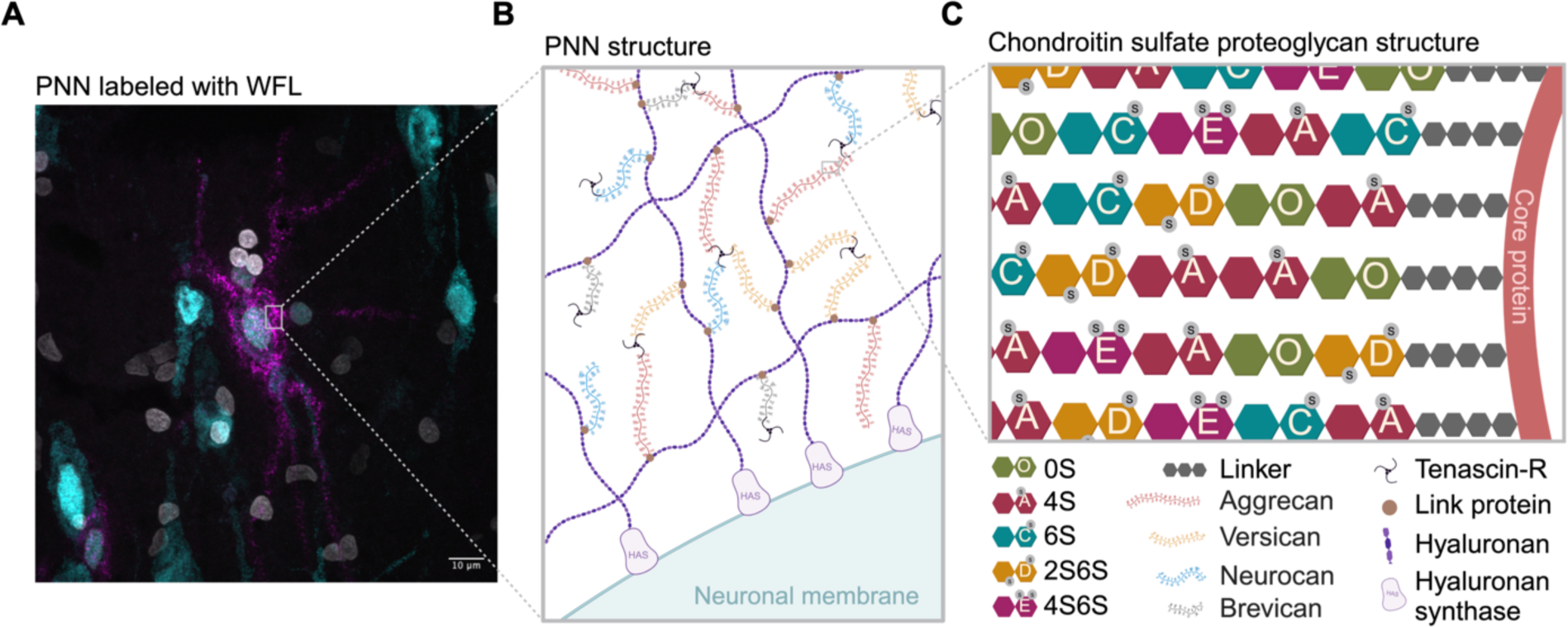
Perineuronal net composition and structure **A** Representative micrograph of a WFL+ PNN (magenta) surrounding the soma and proximal dendrites of a neuron (cyan) in the postmortem human entorhinal cortex. Nuclei (white). **B** PNNs are primarily composed of CSPGs from the lectican family (aggrecan, versican, neurocan and brevican) intricately linked to a hyaluronan backbone synthesized on the cell surface through link proteins. CSPGs are attached to each other through Tenascin-R, resulting in a distinctive net-like structure known as the PNN. **C** CSPGs have varying number of CS-GAG side chains made up of CS-disaccharides that exist in different isomers based on sulfate groups added to either C-4 or C-6 of the GalNAc or C-2 of GlcA including non-sulfated CS-O (0S), mono-sulfated CS-A (4S), -C (6S) and di-sulfated CS-D (2S6S), -E (4S6S). WFL: Wisteria Floribunda Lectin, PNN: perineuronal net, CSPGs: chondroitin sulfate proteoglycans, CS-GAG: chondroitin sulfate glycosaminoglycan, GalNAc: N-acetylgalactosamine, GlcA: D-glucuronic acid, HAS: hyaluronan synthase

PNN CSPGs consist of varying numbers of attached CS glycosaminoglycan (CS-GAG) side chains (Schwartz, N.B. and Domowicz, M.S. 2018). An additional layer of complexity is added by the CS-GAG length as well as the sulfate modifications to each CS disaccharide unit (**Figure 1C**). Sulfate addition can occur on the C-4 and C-6 of the *N*-acetylgalactosamine (GalNAc), and C-2 of D-glucuronic acid (GlcA) that make up the CS disaccharide unit (Djerbal, L., Lortat-Jacob, H., et al. 2017), which form five differentially sulfated CS isomers including non-sulfated (CS-O (0S)), mono-sulfated (CS-A (4S), -C (6S)) and di-sulfated (CS-D (2S6S), -E (4S6S)) variations. The relative abundances of these isomers incorporated into the PNNs constitute a sulfation code (Scarlett, J.M., Hu, S.J., et al. 2022) that relates PNN composition to function (reviewed in (Djerbal, L., Lortat-Jacob, H., et al. 2017)). For example, the ratio of 4-sulfation to 6-sulfation is an important regulator of brain development and aging. In the developing mouse brain (postnatal day (P)0), 27.5% of CS isomers are comprised of 6-sulfation, which is permissive to neurocircuit plasticity and reorganization necessary for neurodevelopment. However, after the end of the critical period (P60), the 6-sulfation drops to 2.3% and instead 87.8% of CS isomers are comprised of 4-sulfation that matures PNN matrices, stabilizes neurocircuitry and limits plasticity of the underlying neurons (Miyata, S., Komatsu, Y., et al. 2012). Extracellular growth factors and proteins, including orthodenticle homeobox protein 2 (Otx2) (Bernard, C. and Prochiantz, A. 2016, Van’t Spijker, H.M., Rowlands, D., et al. 2019), interact with PNN CS-GAG chains and play a crucial role in the proper maturation of PV interneurons and the timing of the critical period of plasticity (Bernard, C. and Prochiantz, A. 2016). For example, PNNs capture Otx2 through protein-glycan binding to the CS-E and CS-D variants, which also triggers maturation of the enmeshed neurons (Beurdeley, M., Spatazza, J., et al. 2012). Notably, overexpression of chondroitin 6-sulfotransferase-1 (C6ST-1) in PV interneurons restricts the formation of stable PNNs and prevents binding of Otx2, resulting in aberrant maturation of PV interneurons and persistent cortical plasticity into adulthood (Miyata, S., Komatsu, Y., et al. 2012). Conversely, specific deletion of the chondroitin 4-sulfotransferase-1 (C4ST-1) gene *Chst11* in mice leads to an augmentation of PNNs in CA2 of the hippocampus (Huang, H., Joffrin, A.M., et al. 2023).

PNNs are also associated with the storage of long-term memories by limiting feedback inhibition from PV cells onto projection neurons (Shi, W., Wei, X., et al. 2019). It is theorized that these extracellular nets preserve spatial information about synapses through the patterning of their holes (Tsien, R.Y. 2013). Fear memories that persist into adulthood are actively preserved by PNNs in the amygdala, as illustrated by a reopening of the critical period and extinction of fear memory in adult mice in which local PNN CS-GAGs are enzymatically degraded using chondroitinaseABC (chABC) (Gogolla, N., Caroni, P., et al. 2009). Due to their protracted development and role in dampening the critical period of brain plasticity in humans and rodents, PNNs have attracted increasing attention in the fields of neurodevelopment (Wen, T.H., Binder, D.K., et al. 2018), neurodegeneration (Lemarchant, S., Wojciechowski, S., et al. 2016) and psychopathology (Browne, C.A., Conant, K., et al. 2022, Carceller, H., Gramuntell, Y., et al. 2023, Pantazopoulos, H. and Berretta, S. 2016).

Early-life stress (ELS) coinciding with the critical period can alter PNN development in the rodent brain (Gildawie, K.R., Honeycutt, J.A., et al. 2020, Guadagno, A., Belliveau, C., et al. 2021, Murthy, S., Kane, G.A., et al. 2019, Santiago, A.N., Lim, K.Y., et al. 2018) but the effects seem to be paradigm-specific, PNN detection method-specific and brain region-specific. We previously reported that PNN density, using WFL to label the CS-GAGs, is significantly increased in the ventromedial prefrontal cortex (vmPFC) of individuals who experienced child abuse compared to controls (Tanti, A., Belliveau, C., et al. 2022). Building upon this, we hypothesize that not only are PNNs increased after child abuse, but their PNN CS-GAG sulfation patterning and consequently their function, are also affected. This study aimed to characterize PNN interregional differences in PNN sulfation coding and labeling patterns across three brain regions: the human vmPFC, entorhinal cortex (EC) and hippocampus (HPC), as well as their corresponding mouse equivalents (mPFC, EC, vHPC). Then, we explored the impact of ELS on these PNN characteristics in both species, recognizing the susceptibility of these brain regions to such influences (Murthy, S. and Gould, E. 2020, Teicher, M.H., Samson, J.A., et al. 2016).

## Results

### PNN sulfation code across the healthy brain

#### Human

To address confounding factors which may contribute to varied findings across brain regions in ELS research, we first used LC-MS/MS to characterize the CS-GAG sulfation code in healthy postmortem human brain tissue across three regions involved in emotion regulation and the threat response: the vmPFC, EC and HPC (Teicher, M.H., Samson, J.A., et al. 2016). In psychiatrically healthy controls, mono-sulfated CS-A (4S) was the most abundant isomer in adult vmPFC (77.6%), EC (76.2%) and HPC (81.2%) compared to CS-C (6S) in the vmPFC (5.6%), EC (8.4%), and HPC (9.5%) (**Figure 2A, Supplementary Table 1**). This finding is consistent with developmental literature suggesting that CS-A replaces CS-C after the closure of the critical window of developmental neuroplasticity (Djerbal, L., Lortat-Jacob, H., et al. 2017, Kitagawa, H., Tsutsumi, K., et al. 1997, Miyata, S., Komatsu, Y., et al. 2012). Meanwhile, di-sulfated CS-D (between 2.0-3.1% of total CS isomers) and CS-E (1.1-1.2% of total CS isomers) were the least abundant in all three brain regions.

**Fig. 2:**
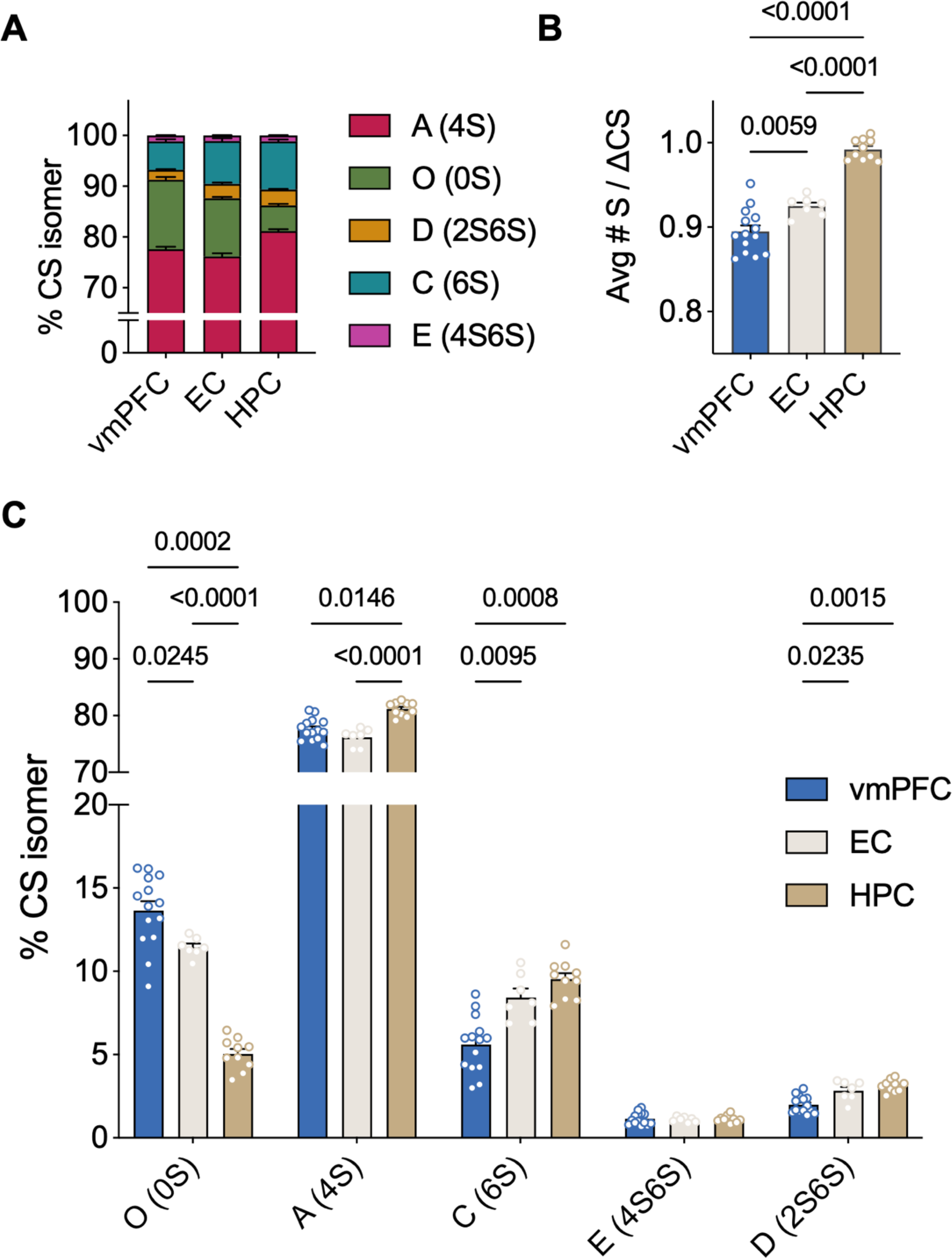
CS-GAG sulfation code varies significantly between brain regions in healthy postmortem human brain **A** Comparison of the relative abundance of each CS isomer in the control vmPFC (N = 14), EC (N = 7) and HPC (N = 10) measured by LC-MS/MS **B** The HPC is the most hypersulfated region on average while the vmPFC is the most hyposulfated region (Welch’s ANOVA: W(2,18) = 101, P < 0.0001, followed by Dunnett’s test) **C** Hypersulfation in the HPC is driven by mono-sulfated CS-A, -C and di-sulfated CS-D, while hyposulfation is driven by high abundance of CS-O in the vmPFC. Each isomer is expressed differently in each brain region except for isomer CS-E (4S6S) (ANOVA: isomer effect: F(2,61) = 20074, P < 0.0001; region effect: F(0.1, 10) = 5.478e-20, P > 0.9999; isomer x region: F(2,43) = 54.23, P <0.0001, followed by Tukey’s test). Avg#S/7CS: average number of sulfates per chondroitin sulfate disaccharide, CS-GAG: chondroitin sulfate glycosaminoglycan, vmPFC: ventromedial prefrontal cortex, EC: entorhinal cortex, HPC: hippocampus, LC-MS/MS: liquid chromatography tandem mass spectrometry, ANOVA: analysis of variance.

When testing for differences in sulfation patterning between human brain regions, we found significant variations between cortical and deep brain structures. The HPC (0.99 Avg#S/ΔCS) exhibited an increase in hypersulfation compared to the EC (0.92 Avg#S/ΔCS) and vmPFC (0.90 Avg#S/ΔCS) (**Figure 2B**). The hypersulfation observed in the HPC is driven by both the high abundance of mono- and di-sulfated isomers (CS-A, -C, -D), and decreased abundance of non-sulfated isomer (CS-O), specific to this brain region (**Figure 2C**). Meanwhile, the hyposulfation observed in the vmPFC was dependent on the decreased abundance of sulfated CS-C and CS-D and increase in non-sulfated CS-O (**Figure 2C**). In comparison, the EC presented an intermediate Avg#S/ι1CS and abundance of non-sulfated, mono-sulfated and di-sulfated isomers when compared to the HPC and vmPFC (**Figure 2B-C**).

#### Mouse

To determine whether these region-specific differences in PNN CS-GAG sulfation profiling is conserved across species, we next determined the relative abundance of each CS isomers using LC-MS/MS in fixed, wild type mouse brain. Similar to humans (**Figure 2**), control mice showed CS-A as the most abundant isomer in the mPFC (70%), EC (73.9%), and vHPC (79.6%) (**Figure 3A, Supplementary Table 2**), while di-sulfated CS-E was the least abundant with 1.4% in the mPFC, 0.9% in the EC and 0.7% in the vHPC.

**Fig. 3:**
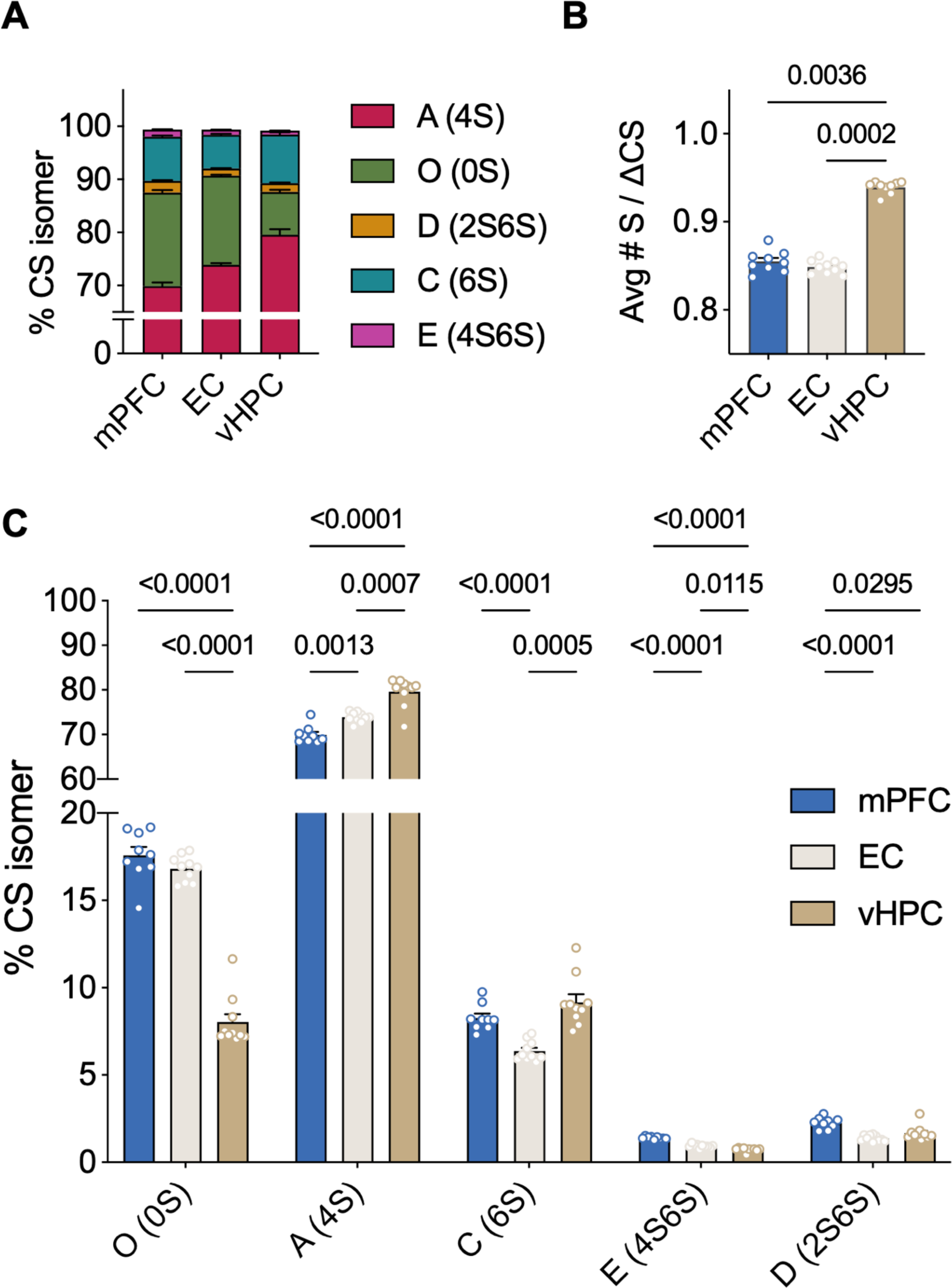
CS-GAG sulfation code is generally conserved between species in the healthy brain **A** Comparison of the relative abundance of each CS isomer in the control mPFC, EC and vHPC of mouse (N = 10) brain measured by LC-MS/MS **B** Like human brain, the mouse vHPC is hypersulfated compared to both the EC and mPFC, however, in this species the mPFC and EC do not differ in average sulfation (Kruskal-Wallis test: P = 0.0001, followed by Dunn’s test) **C** All CS-isomers are expressed in significantly different abundances across the healthy mouse brain (ANOVA: isomer effect: F(1,11) = 13984, P < 0.0001; region effect: F(1, 12) = 0.01896, P = 0.940; isomer x region : F(2, 16) = 97.49, P < 0.0001, followed by Tukey’s test). Avg#S/ΔCS: average number of sulfates per chondroitin sulfate disaccharide, CS-GAG: chondroitin sulfate glycosaminoglycan, LC-MS/MS: liquid chromatography tandem mass spectrometry, mPFC: medial prefrontal cortex, EC: entorhinal cortex, vHPC: ventral hippocampus, ANOVA: analysis of variance.

We also observed parallel differences in CS-GAG sulfation patterning between mouse brain regions, whereas the vHPC exhibited CS-GAG hypersulfation (0.94 Avg#S/ι1CS), both mPFC (0.85 Avg#S/ι1CS) and EC (0.85 Avg#S/ι1CS) were hyposulfated (**Figure 3B**). vHPC hypersulfation appeared to be driven by the high abundance of mono-sulfated isomers CS-A and CS-C and low abundance of non-sulfated CS-O (**Figure 3C**) compared to EC and mPFC. Unlike the human samples, the mPFC (17.6% of CS isomers) and EC (16.8% of CS isomers) exhibited similar CS-O sulfation levels.

### Labeling patterns of PNNs across the healthy brain

#### Human

To characterize the labeling pattern of PNNs in relation to sulfation changes, tissues were co-labeled with both ACAN (PNN core protein) and WFL (PNN CS-GAGs). Matching previous reports, healthy human brain showed strong PNN labeling heterogeneity, including nets that appeared deglycosylated (WFL−/ACAN+) and those appearing glycosylated (WFL+/ACAN− and WFL+/ACAN+) (Härtig, W., Meinicke, A., et al. 2022, Matthews, R.T., Kelly, G.M., et al. 2002, Miyata, S., Nadanaka, S., et al. 2018, Scarlett, J.M., Hu, S.J., et al. 2022, Ueno, H., Fujii, K., et al. 2018) (**Figure 4A-B**). ACAN immunoreactivity was found to be associated with extracellular PNN formations (**Figure 4B yellow arrow**) but also within the cytoplasm of some cells (**Figure 4B cyan arrow**), as previously reported in neurons (Domowicz, M., Mangoura, D., et al. 2000) as well as glia (Asher, R.A., Scheibe, R.J., et al. 1995). Therefore, only ACAN labeling appearing as extracellular PNNs was included in further analyses (**Figure 4B**). We found that PNN heterogeneity varied significantly between brain regions (**Supplementary Table 3**), whereas the vmPFC exhibited the greatest amount of double labeled PNNs (48.3% WFL+/ACAN+) compared to the HPC (20.1%) and EC (12.7%) (**Figure 4C**). The EC presented the most WFL+/ACAN− labeling (66.8% of total PNNs) compared to HPC (25.7%) and vmPFC (28.3%), and the HPC exhibited the greatest amount of deglycosylated ACAN+ PNNs (54.2% WFL−/ACAN+) compared to the EC (20.5%) and vmPFC (23.4%) suggesting additional PNN core proteins may prevail in this region.

**Fig. 4:**
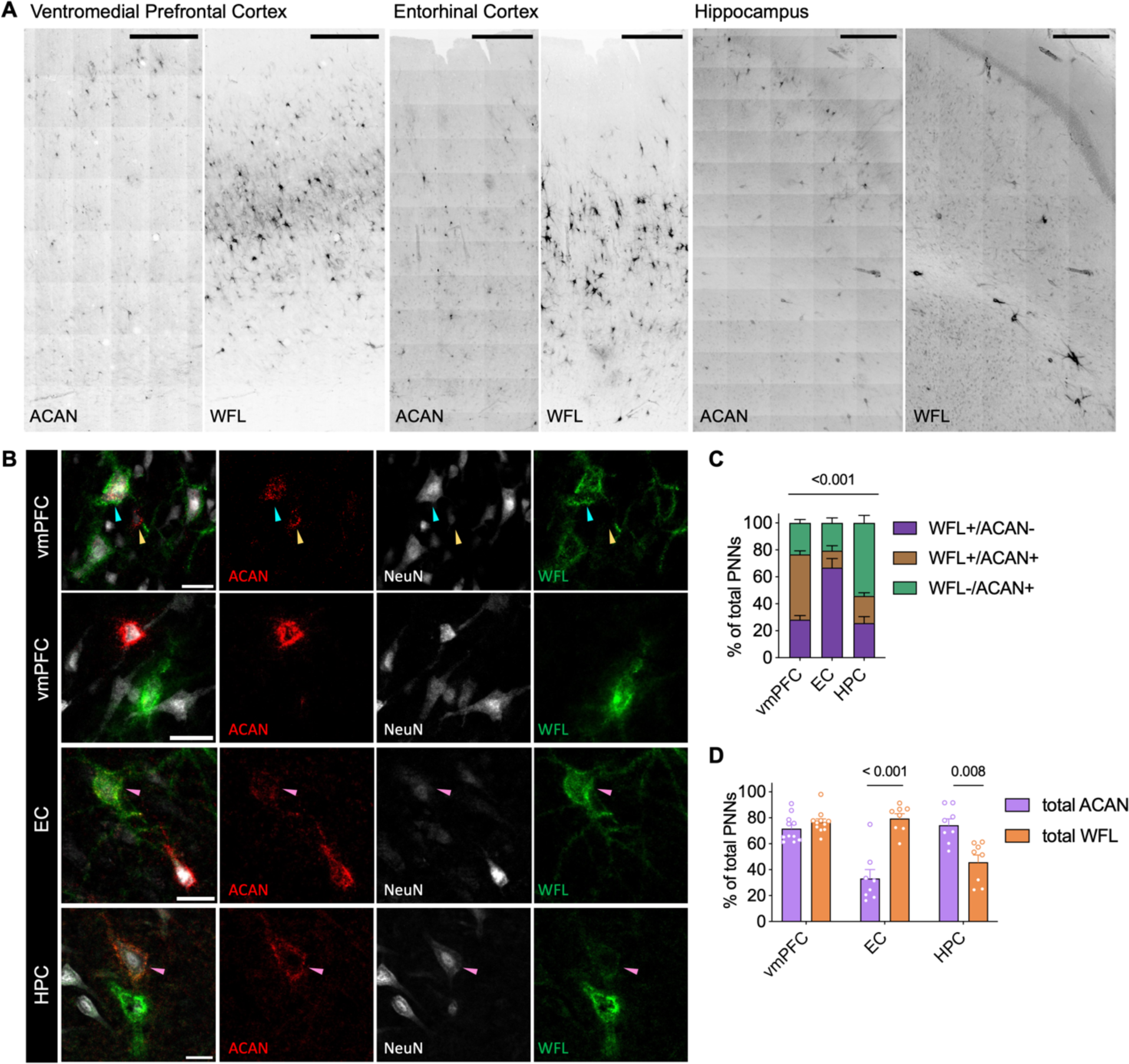
Immunohistochemical staining of healthy human brain reveals heterogeneity in perineuronal net labeling that differs between brain regions **A** Representative micrographs of the distribution of either ACAN or WFL labeling across the human vmPFC (N = 11), EC (N = 8) and HPC (N = 8). Imaged at 20X on Evident Scientific VS120 Slide Scanner, scale bar 500µm. **B** PNNs were manually classified as either single (WFL+/ACAN− or WFL−/ACAN+) or double (WFL+/ACAN+, pink arrows) labeled. ACAN staining can be found either intracellularly (100% colocalization with NeuN, cyan arrow) or in the distinctive pattern of an extracellular PNN (surrounding NeuN, yellow arrow), only the latter was considered as an ACAN+ PNN in this study (yellow arrow). Imaged at 40X on Evident Scientific FV1200 Confocal, scale bar 25µm. **C** Comparison of the labeling patterning between the different brain regions investigated (ANCOVA with age as covariate: labeling x region: F(3,46) = 25.3, p < 0.001). **D** Total ACAN (WFL+/ACAN+ and WFL−/ACAN+) and total WFL (WFL+/ACAN− and WFL+/ACAN+) labeling can be found in different amounts depending on the brain region (ANCOVA with age as covariate: total labeling x region: F(2,23) = 19.5, p < 0.001). Within the vmPFC the two markers label similarly, while WFL prevails in the EC and ACAN predominates in the HPC. WFL: Wisteria Floribunda Lectin, ACAN: anti-aggrecan core protein, NeuN: anti-neuronal nuclei, vmPFC: ventromedial prefrontal cortex, EC: entorhinal cortex, HPC: hippocampus, ANCOVA: analysis of co-variance.

We then hypothesized that differences in PNN heterogeneity may be related to differences in PNN CS-GAG sulfation patterning unique to each brain region. By examining the total labeling of a marker (total ACAN labeling includes both the WFL−/ACAN+ and WFL+/ACAN+; total WFL labeling includes WFL+/ACAN− and WFL+/ACAN+ PNNs), we observed large differences in total labeling between brain regions (**Figure 4D**). Compared to the vmPFC that showed equal abundance of ACAN (71.7%) vs WFL (76.6%) total PNN labeling, the EC exhibited a significantly higher total amount of WFL (79.4%) labeled PNNs compared to ACAN (33.2%) labeled PNNs. In contrast, the HPC exhibited a significantly higher amount of ACAN (74.3%) labeled PNNs compared to WFL (45.8%) labeled PNNs (**Figure 4D**).

#### Mouse

To investigate whether labeling pattern of PNNs is conserved across species, healthy mouse brain was also co-labeled with WFL and ACAN (**Figure 5A**). Surprisingly, the labeling pattern of PNNs is highly dissimilar between species. The specificity of the ACAN antibody used in mouse was demonstrated by intracellular ACAN immunoreactivity (as observed in human samples and not considered for analysis, **Figure 5B, cyan arrow**) as well as by the abundance of ACAN+ PNNs in the adjacent brain regions within the same coronal sections (**Figure 5A**). In the mouse mPFC and EC, 94.7% and 92.3%, of all PNNs were labeled as WFL+/ACAN− (**Figure 5C, Supplementary Table 4**). Nearly all other PNNs in these brain areas were found to be double positive for WFL+/ACAN+ (5.5-7.1%), while very few PNNs in the mPFC and EC labeled as WFL−/ACAN+ (0-0.6%, **Figure 5C**). In contrast, the vHPC displayed more heterogeneity in PNN labeling with all three populations being present: 12.5% WFL−/ACAN+, 39.5% WFL+/ACAN− and 47.9% WFL+/ACAN+ (**Figure 5B**). Once again, the mPFC and EC appeared to display much higher levels of glycosylation than the vHPC, as seen in humans.

**Fig. 5:**
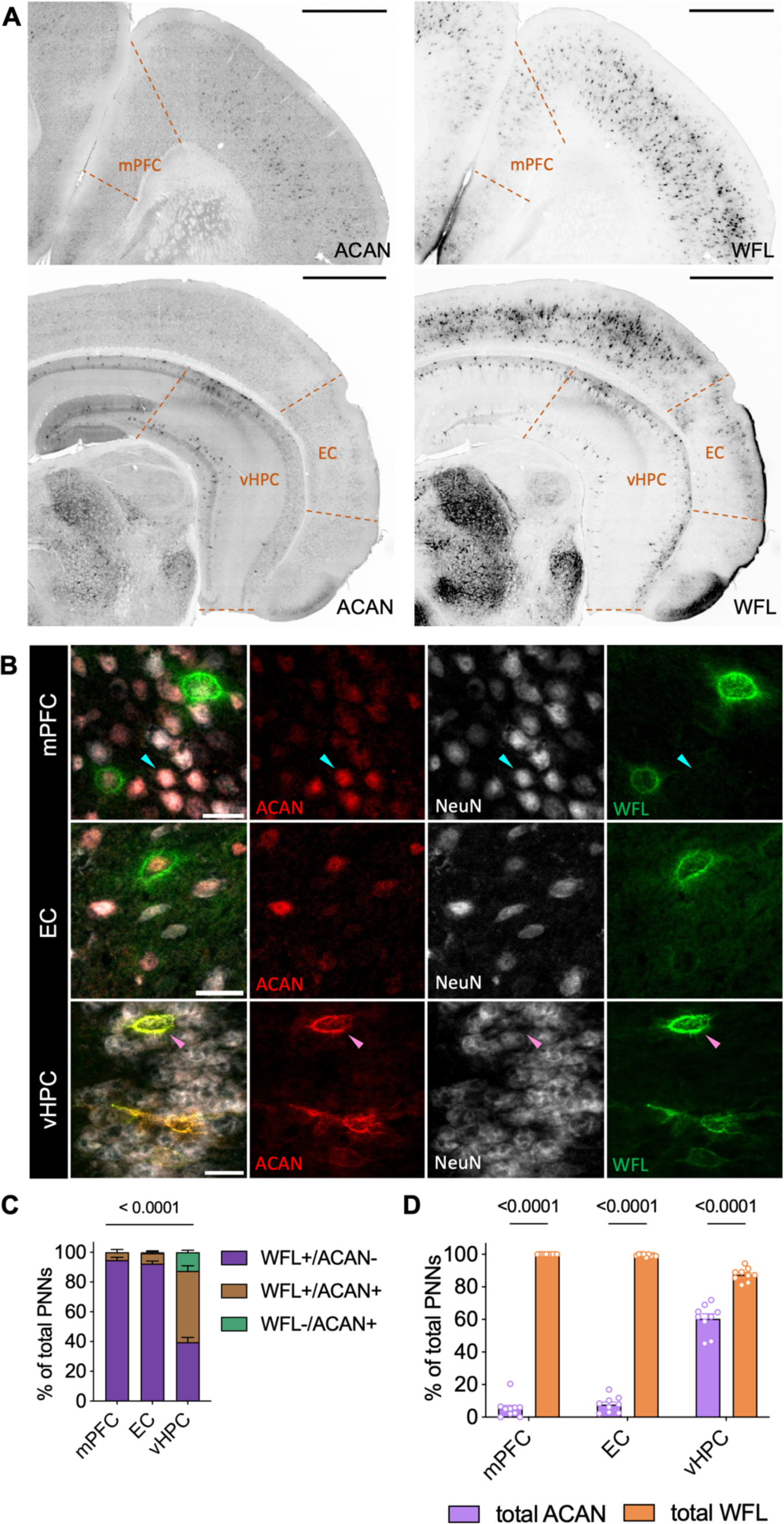
Perineuronal net labeling is highly dissimilar between species and brain region **A** Representative micrographs of the distribution of either ACAN or WFL labeling across control mouse mPFC, EC and vHPC (N = 10). Imaged at 20X on Evident Scientific VS120 Slide Scanner, scale bar 500µm. **B** PNNs were manually classified as either single (WFL+/ACAN− or WFL−/ACAN+) or double (WFL+/ACAN+, pink arrow) labeled. As in the human samples, ACAN staining can be found either intracellularly (100% colocalization with NeuN, cyan arrow) or in the distinctive pattern of an extracellular PNN (surrounding NeuN, pink arrow), only the latter was considered as an ACAN+ PNN in this study. Imaged at 40X on Evident Scientific FV1200 Confocal, scale bar 25µm. **C** PNN labeling pattern varies significantly by brain region (ANOVA: labeling x region F(4,50) = 140.4, p < 0.0001). Unlike the human samples, barely any PNNs were labeled solely by WFL−/ACAN+ in control mouse mPFC and EC. **D** Examination of total WFL or ACAN labeling within brain regions reveals that WFL labels significantly more PNNs in all brain regions examined (ANOVA: total labeling x region F(2,32) = 330.2, p < 0.0001). WFL: Wisteria Floribunda Lectin, ACAN: anti-aggrecan core protein, mPFC: medial prefrontal cortex, EC: entorhinal cortex, vHPC: ventral hippocampus.

When comparing the total labeling of ACAN (WFL−/ACAN+, WFL+/ACAN+) or WFL (WFL+/ACAN+, WFL+/ACAN−) between regions, we observed a similar labeling pattern of WFL across the brain in mice and humans, but ACAN was differentially dispersed (**Figure 5D**). In the mPFC and EC, 99.4-100% of PNNs were labeled with WFL, while in the vHPC only 87.5% of PNNs were labeled. In contrast, the vHPC exhibited 60.5% of PNNs labeled with ACAN while the mPFC and EC only had 5.4-7.7% of PNNs stained with ACAN respectively.

### ELS influences PNN abundance but not sulfation code

#### Human

Although we previously reported an increase in density of WFL+ PNNs in the vmPFC of individuals who died by suicide during a depressive episode with a history of child abuse (Tanti, A., Belliveau, C., et al. 2022), there was no significant difference in the relative abundance of CS isomers associated with suicide (DS) or abuse (DS-CA) in any of the three regions analysed compared to controls (CTRL) (**Figure 6A-C**). Additionally, the labeling pattern landscape did not vary with abuse throughout the brain (**Figure 6D-F**).

**Fig. 6:**
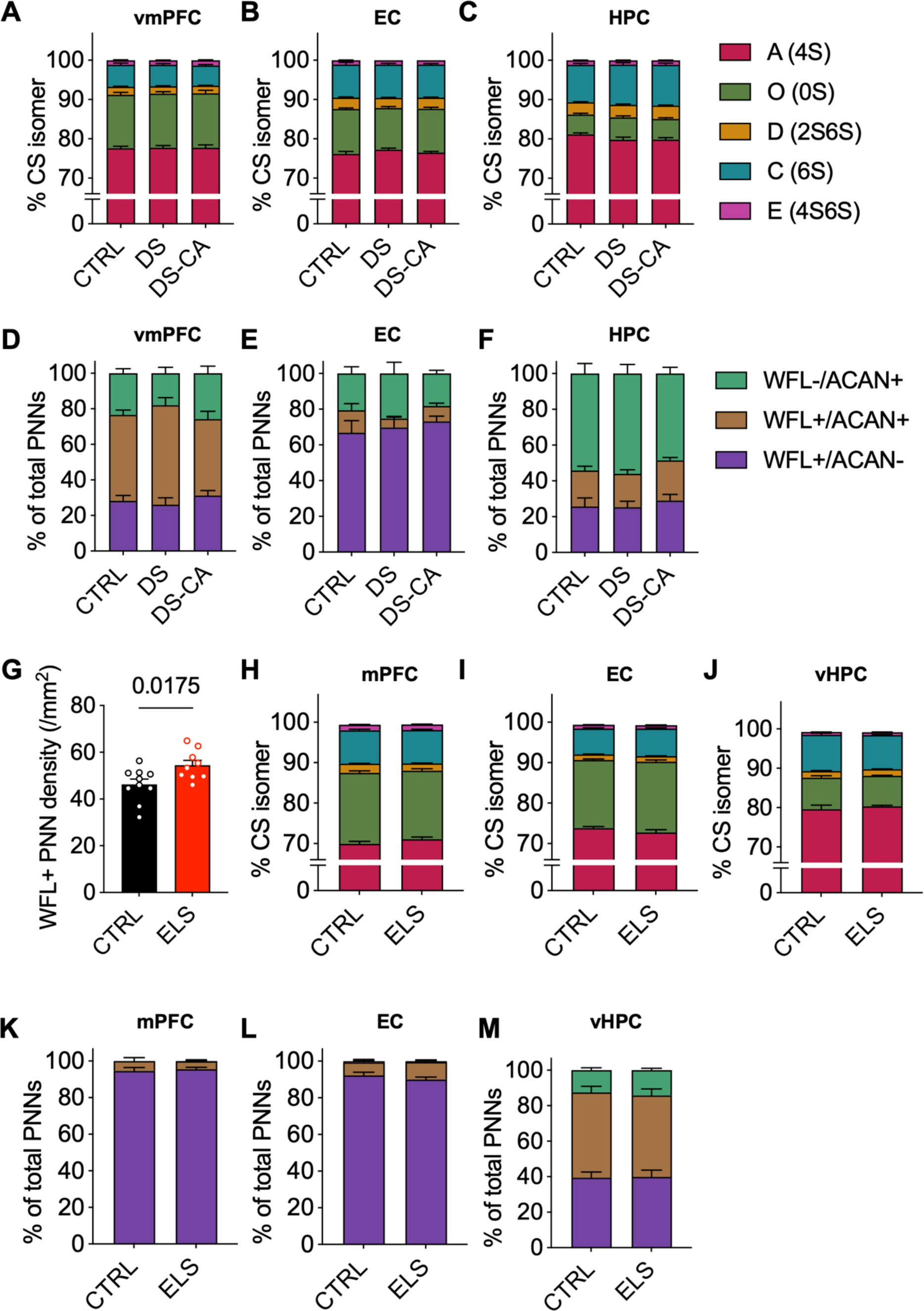
Perineuronal net sulfation code and labeling pattern are preserved after the experience of early-life stress in both human and mouse brain **A-C** Within each human brain region CS-GAG sulfation code is unaffected by depression, suicide (DS) and child abuse (DS-CA) when compared to controls (CTRL) (ANOVA: vmPFC group x isomer: F(3,67) = 0.132, P =0.929; EC group x isomer: F(3,13) = 1.448, P = 0.275; HPC group x isomer: F(2,6) = 1.210, P = 0.372) **D-F** Although we previously reported a significant increase in WFL+ PNNs in the vmPFC, overall labeling pattern is conserved between groups in all brain regions examined (ANCOVA with age as covariate: vmPFC group x labeling: F(4,50) = 2.104, P = 0.094; EC group x labeling: F(3,38) = 0.471, P = 0.65; HPC group x labeling: F(3,47) = 0.152, P = 0.961) **G** Limited bedding and nesting from P2-9 is associated with a significant increase in WFL+ PNNS in the mPFC of adult mice (N = 9) compared to control (N =10) (Welch’s t-test: P = 0.0175) **H-J** Within each mouse brain region CS-GAG sulfation code is unaffected by early-life stress (ANOVA: mPFC group x isomer: F(1,21) = 1.761, P = 0.200; EC group x isomer: F(1,9) = 5.029, P = 0.464 although does not pass Bonferroni’s multiple comparisons test; vHPC group x isomer: F(1,22) = 0.683, P = 0.424) **K-M** Although we report an increase in WFL+ PNNs in the mPFC of mice, the overall landscape of PNN labeling by WFL and ACAN is resilient to ELS (ANOVA: mPFC group x labeling: F(2,34) = 0.248, P = 0.782; EC group x labeling: F(2,32) = 1.33, P = 0.279; vHPC group x labeling: F(2, 32) = 0.140, P = 0.870). CTRL: psychiatrically healthy control, DS: depressed suicide, DS-CA: depressed suicide with a history of child abuse, CS-GAG: chondroitin sulfate glycosaminoglycan, WFL: Wisteria Floribunda Lectin, ACAN: anti-aggrecan core protein, vmPFC: ventromedial prefrontal cortex, EC: entorhinal cortex, HPC: hippocampus, ANCOVA: analysis of covariance, mPFC: medial prefrontal cortex, EC: entorhinal cortex, vHPC: ventral hippocampus, ANOVA: analysis of variance.

#### Mouse

Initially, to validate our mouse model of ELS, we examined the density of WFL+ PNNs in the mPFC. Replicating our previously published human data (Tanti, A., Belliveau, C., et al. 2022) we found that there are significantly more WFL+ PNNs in the mPFC of ELS mice (N = 9) compared to CTRL mice (N = 10) (**Figure 6G**). Although, PNN CS-GAG abundance appeared elevated, ELS did not influence the underlying CS-GAG sulfation code (**Figure 6H-J**) or labeling pattern (**Figure 6K-M**) in any of the three mouse brain regions investigated. These results suggest that ELS only has a long-lasting effect on PNN CS-GAG abundance and not composition, an observation we found conserved between species.

## Discussion

We observed and characterized for the first time vastly different relative abundances of CS disaccharide isomers and PNN labeling patterns between brain regions across species, shedding new translational light on the intricate foundations of PNN composition. We initially predicted there would be a higher amount of double labeled PNNs (WFL+/ACAN+) in all brain regions across species based on evidence suggesting that ACAN is necessary for PNN formation *in vitro* (Giamanco, K.A., Morawski, M., et al. 2010) and in vivo (Rowlands, D., Lensjø, K.K., et al. 2018). Although most studies investigating changes in PNN abundance are conducted using WFL as a universal PNN marker and a widely accepted histochemical visualization method, WFL was recently shown to preferentially bind to non-sulfated CS disaccharide (CS-O) (Nadanaka, S., Miyata, S., et al. 2020). Although we did not observe a significant correlation between labeling pattern and sulfation score *per se* (**Supplementary Figure 1**), we did find that brain regions with the highest levels of non-sulfated CS-O also exhibited the strongest PNN labeling with WFL. In both healthy human and mouse brain, the (v)mPFC and EC had the most glycosylation (total WFL labeling) and the most CS-O. In comparison, the (v)HPC had the least glycosylation and lowest percentages of CS-O. These findings are consistent with previous research showing increased CS-O content in the somatosensory cortex correlated to elevated WFL+ PNN labeling in P45 mice when compared to the adjacent dorsal hippocampus (Scarlett, J.M., Hu, S.J., et al. 2022). Interestingly, the hippocampus is also the brain region with the highest amount of CS-A which has been shown to be useful for creating a healthy niche for neural stem cells through its ability to bind basic fibroblast growth factor (Djerbal, L., Lortat-Jacob, H., et al. 2017). Overall, these findings highlight the limitations of using WFL as a pan-PNN labeling marker.

This study emphasizes the importance of using multiple markers in order to accurately represent the degree of regional PNN diversity throughout the brain. It was originally thought that PNNs would double label when combining WFL labeling and immunohistochemistry with anti-CSPG antibodies (Härtig, W., Brauer, K., et al. 1994). However, more recent evidence suggests that WFL does not always co-label PNNs detected with anti-CSPG antibodies (Härtig, W., Meinicke, A., et al. 2022, Matthews, R.T., Kelly, G.M., et al. 2002, Miyata, S., Nadanaka, S., et al. 2018, Scarlett, J.M., Hu, S.J., et al. 2022, Ueno, H., Fujii, K., et al. 2018), including anti-aggrecan (ACAN). Moreover, rodent literature shows that different CSPG antibodies bind to different portions of the large proteoglycan such that three different antibodies for ACAN (Cat-301, Cat-315 and Cat-316) bind selectively to this target at unique sites. Nevertheless, their labeling pattern is completely different depending on the brain region and glycosylation pattern of the CSPG (Matthews, R.T., Kelly, G.M., et al. 2002). The adult human cortex, specifically Brodmann area 6, expresses all forms of ACAN isoforms (Virgintino, D., Perissinotto, D., et al. 2009) albeit to different extent throughout the cortical neuronal layers and enmesh different cell types. In our study, Cat-301, which labels the core protein itself, was the anti-ACAN antibody used to label human brain samples. In contrast, Cat-316 binds selectively to non-sulfated CS-GAG on ACAN and labels more similarly to WFL (Matthews, R.T., Kelly, G.M., et al. 2002), thus producing a different landscape of ACAN PNN labeling throughout the brain. The antibody used in the mouse brain for this study also binds to the core protein itself, as apparent by the molecular weight reported by the manufacturer (**Table II**).

The unique composition of PNNs in different brain regions and species, as seen here and by others (Lensjø, K.K., Christensen, A.C., et al. 2017, Matthews, R.T., Kelly, G.M., et al. 2002), suggests region-specific roles in regulating plasticity and memory processes. Literature indicates that PNN formation may be dependent on neuronal activity (Devienne, G., Picaud, S., et al. 2021, Kind, P.C., Sengpiel, F., et al. 2012, McRae, P.A., Baranov, E., et al. 2010, Rowlands, D., Lensjø, K.K., et al. 2018) implying that regions with more labeling are highly active compared to regions with less labeling. Interestingly, this phenomenon has been observed using both WFL and ACAN to visualize the nets, although none of the indicated studies examined both markers at the same time. Given PNN heterogeneity in both CS core proteins and associated CS-GAG sulfation patterns, future studies should explore their composition at the single-cell level, as this should shed light on the functional consequences of varied sulfation patterns on different neuronal subtypes across brain regions and species.

The present study also investigated the long-lasting impact of early-life adversity on PNN heterogeneity and composition in the (v)mPFC, EC and (v)HPC of adult human and mouse brain. Using an ELS mouse model of limited bedding and nesting from P2-9 (Reemst, K., Kracht, L., et al. 2022), we found that WFL+ PNNs are increased in the mPFC of P70 male and female mice compared to controls (**Figure 6**). Moreover, our findings elucidate that PNN sulfation code and labeling pattern are undisturbed in adult human and mouse brains that have experienced early-life adversity. We cannot exclude, however, that this code was altered earlier in life during and/or shortly after the ELS but then restored thereafter. Biochemical composition changes of PNNs are not solely restricted to the critical period of plasticity early in life. For example, glia are becoming increasingly involved in the production, maintenance and degradation of cerebral PNNs (Crapser, J.D., Arreola, M.A., et al. 2021) in adult mice (Venturino, A., Schulz, R., et al. 2021) and humans (Crapser, J.D., Spangenberg, E.E., et al. 2020) in health and disease. Glia are also known to produce and secrete C4ST-1 (Bhattacharyya, S., Zhang, X., et al. 2015) and C6ST-1 (Properzi, F., Carulli, D., et al. 2005), main regulators of CS-GAG sulfation. It will be important to further investigate how cerebral PNNs are affected by early-life adversity, given the rich body of research highlighting the long-lasting impact of ELS on WFL+ PNNs across various paradigms and brain regions in different species (Gildawie, K.R., Honeycutt, J.A., et al. 2020, Guadagno, A., Belliveau, C., et al. 2021, Murthy, S., Kane, G.A., et al. 2019, Santiago, A.N., Lim, K.Y., et al. 2018, Tanti, A., Belliveau, C., et al. 2022).

This study is not without limitations. Firstly, we examined bulk tissue for LC-MS/MS and not individual PNNs and acknowledge that both interstitial and PNN CS-GAGs may have been included in our analyses. Seeing as each PNN exists on a spectrum: being diffuse to being a highly organized structure, the LC-MS/MS protocol used herein assesses the relative abundance of CS isomers isolated from all matrix subtypes. We previously reported that the CS-GAG sulfation patterns reported utilizing PNN fractionation are reproducible using this LC-MS/MS protocol (Alonge, K.M., Logsdon, A.F., et al. 2019), supporting the idea that the signatures reported here as the sulfation code are driven predominantly by PNNs, whether diffuse or highly structured. Second, the binding of the anti-aggrecan antibody selected in this study (Cat-301) can be inhibited by the presence of CS-GAG chains attached to the core protein (Matthews, R.T., Kelly, G.M., et al. 2002). Future studies examining the landscape of PNN labeling using various anti-aggrecan antibodies should explore tissue pre-treatment with chABC. This pre-treatment has the potential to uncover more of the core protein in ACAN+ PNNs, facilitating their further characterization. However, it is essential to note that chABC pre-treatment would eliminate CS-GAG labeling by WFL. Consequently, we opted not to use chABC pre-treatment, as our primary objective was to analyze the colocalization or lack thereof, between WFL and ACAN. We also acknowledge that more detailed, multi-point longitudinal studies in animal models of ELS should be conducted to further investigate temporal changes in sulfation code and resultant labeling.

## Materials and Methods

### Human post-mortem brain samples

Brain samples from a total of 81 individuals were obtained from the Douglas-Bell Canada Brain Bank (DBCBB; Montreal, Canada). In collaboration with Quebec’s Coroner and the McGill Group for Suicide Studies, informed consent was obtained from the next of kin. Standardized psychological autopsies, medical records and Coroner reports were used to define cases and controls. Any presence of suspected neurodevelopmental, neurodegenerative, or neurological disorders in clinical files were basis for exclusion. In addition, proxy-based interviews with one or more informants that were closely acquainted with the deceased were used to create clinical vignettes with detailed medical and personal histories. Characterization of early-life histories were based on adapted Childhood Experience of Care and Abuse (CECA) interviews addressing the experiences of sexual and physical abuse as well as neglect (Bifulco, A., Brown, G.W., et al. 1994, Lutz, P.E., Tanti, A., et al. 2017). Reports of non-random major physical and/or sexual abuse during the first 15 years of life with a score of 1 or 2 on the CECA were considered as severe child abuse. A panel of clinicians reviewed and assessed all clinical vignettes to generate Diagnostic and Statistical Manual of Mental Disorders (DSM-IV) diagnostic criteria and classify the deceased as a case or control. Toxicology reports and medication prescriptions were also obtained. Three groups were compared in this study consisting of healthy individuals having died suddenly (CTRL; n=19), and depressed suicides with (DS-CA; n=33) or without (DS; n=30) a history of severe child abuse (characteristics detailed in **Table I**).

**Table I:** Group characteristics.

| **Table I: Group characteristics** |  |  |  |
| --- | --- | --- | --- |
|  | **CTRL** | **DS** | **DS-CA** |
| **N = 81** | 18 | 30 | 33 |
| **Axis 1 diagnosis** | 0 | MDD (25); DD-NOS (5) | MDD (26); DD-NOS (7) |
| **Age (years) (P = 0.06)** | 48.1 ± 20.2 | 50.4 ± 12.5 | 41.5 ± 14.6 |
| **Sex (M/F)** | 13/5 | 25/5 | 22/11 |
| **PMI (h) (P = 0.14)** | 41.1 ± 25.0 | 54.0 ± 29.0 | 56.9 ± 27.7 |
| **Tissue pH (P = 0.99)** | 6.5 ± 0.3 | 6.5 ± 0.4 | 6.5 ± 0.4 |
| **Substance dependence** | 0 | 8 | 8 |
| **Last 3 months medication** | Sedative hypnotics (1) | Classic antidepressants (4); Benzodiazepines (9); SNRI (4); SSRI (6); Antipsychotic (3); TCA (2); Antimanic (1); Opioid (1) | Classic antidepressants (2); Benzodiazepines (6); SNRI (3); SSRI (7); Antipsychotic (3); TCA (1); Antimanic (1); Opioid (2); NASSA (2); SARI (1); Anticonvulsant (2); GABA analog (1) |

### Human tissue dissections and preparation

Expert staff at the DBCBB dissected 0.5cm-thick coronal sections from isopentane snap-frozen brain samples with the guidance of the human brain atlas (Juergen Mai, M.M., George Paxinos 2016). vmPFC samples were dissected from coronal sections equivalent to plate 3 (approximately −48mm from the center of the anterior commissure) of this atlas. Anterior HPC samples were obtained from sections equivalent to plate 37 of the atlas, by dissecting around hippocampal CA1-CA3, then between the EC and the parahippocampal gyrus. Samples were then fixed in 10% formalin for 24 h, followed by storage in 30% sucrose at 4°C until they sank. Tissue blocks were snap-frozen in isopentane at −35°C before being stored at −80°C until they were sectioned on a cryostat. 40μm-thick free-floating serial sections were stored in cryoprotectant at −20°C until use.

### Early-life stress mouse model

Animal housing and experiments followed the Canadian Council on Animal Care (CCAC) guidelines, procedures were approved by the Animal Care Committee of the Douglas Research Center (Protocol number 5570) and all experiments were conducted in accordance with ARRIVE guidelines (Percie du Sert, N., Hurst, V., et al. 2020). 8-week-old male and female C57BL/6J mice were purchased from Charles Rivers (Wilmington, MA, USA). Mice were kept at the Douglas animal facility for at least a week before being bred by housing a single female with one male for one week. Females were monitored daily for the birth of pups. ELS was induced using the limited bedding and nesting paradigm, which causes fragmented maternal care from P2 to P9, resulting in chronic stress in pups (Reemst, K., Kracht, L., et al. 2022, Rice, C.J., Sandman, C.A., et al. 2008). Briefly, at P2, in the ELS group, mother and litters were moved to cages with a small amount of corn husk bedding at the bottom and half of the standard amount of nesting material (2.5 x 5 cm piece of cotton nesting material) on top of a stainless-steel mesh which was placed 1 cm above the cage floor. Animals were left undisturbed until P9, when they were moved back to cages with regular housing conditions. CTRL cages were equipped with standard amounts of bedding and nesting material (5 x 5cm). Mice had access to standard chow and water *ad libitum* and were kept under a 12:12 h light-dark cycle. Animals were weaned at P21 and sacrificed at P70. Both male (CTRL = 5, ELS = 9) and female (CTRL = 5, ELS =5) mice were used in this study and no sex differences were observed.

### Mouse brain preparation

At P70, mice were deeply anesthetized with an intraperitoneal injection of ketamine/xylazine (100/10 mg/kg, Sigma-Aldrich, St. Louis, MO, USA) and perfused transcardially with phosphate buffered saline (PBS) followed by 4% paraformaldehyde (PFA, pH 7.4, dissolved in PBS). Brains were then extracted and post-fixed overnight in 4% PFA at 4 °C and subsequently cryoprotected in 30% sucrose dissolved in PBS for 48 h at 4 °C. Sunken brains were snap-frozen in isopentane at −35°C. Lastly, 35μm-thick serial coronal brain sections were collected free-floating and stored in cryoprotectant at −20°C until use.

### Liquid chromatography tandem mass spectrometry

Six to ten serial sections from human vmPFC (CTRL = 14, DS = 16, DS-CA = 12), EC (CTRL = 7, DS = 21, DS-CA = 22), HPC (CTRL = 10, DS = 19, DS-CA = 20) or mouse (CTRL = 10, ELS = 9) mPFC, EC, vHPC samples were processed for liquid chromatography tandem mass spectrometry (LC-MS/MS) as previously described (Alonge, K.M., Logsdon, A.F., et al. 2019, Logsdon, A.F., Francis, K.L., et al. 2022). Briefly, samples were shipped from Montreal to Seattle in PBS + 0.02% sodium azide at 4°C. After manual dissections of the brain regions of interest (ROI), sections were washed thrice in Optima LC/MS-grade water and once in 50mM ammonium bicarbonate (pH 7.6) at room temperature (RT). Tissues were incubated with 50mU/mL of chABC (Sigma-Aldrich, C3667, Burlington, Massachusetts, USA) for 24 h at 37°C. The supernatant was filtered and lyophilized, then reconstituted in LC/MS-grade water for LC-MS/MS analysis.

### LC-MS/MS + MRM quantification of isolated chondroitin sulfate disaccharides

Human and mouse CS isomers were subjected to quantification using a triple quadrupole (TQ) mass spectrometer equipped with an electrospray ion (ESI) source (Waters Xevo TQ-S) and operated in negative mode ionization as previously described (Alonge, K.M., Logsdon, A.F., et al. 2019, Logsdon, A.F., Francis, K.L., et al. 2022, Scarlett, J.M., Hu, S.J., et al. 2022). Briefly, CS disaccharides were resolved on porous graphite column (Hypercarb column; 2.1 × 50 mm, 3 μm; Thermo Fisher Scientific, Bothell, WA, USA) and LC-MS/MS + MRM were performed using a Waters Acquity I-class ultra-performance liquid chromatographic system (UPLC) using the following MRM channels: CS-A (4S), *m/z* 458 > 300; CS-C (6S), *m/z* 458 > 282; CS-D (2S6S) and CS-E (4S6S), *m/z* 268 > 282; CS-O (0S), *m/z* 378 > 175. MassLynx software version 4.1 (Waters) was used to acquire and quantify all data using a modified peak area normalization function (Alonge, K.M., Logsdon, A.F., et al. 2019). Each CS disaccharide is shown as the relative percentage of all CS isomers within the sample. CS sulfation was computed using the weighed mean formula for the average number of sulfates (Avg#S) per CS disaccharide (ΔCS):

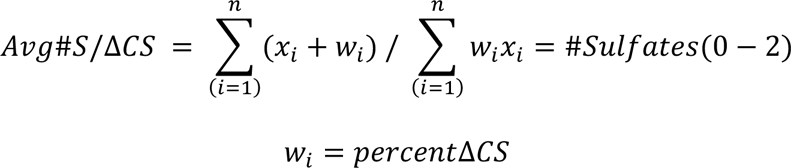

### Immunolabelings, WFL staining, and image analysis

Free-floating sections of human or mouse brain were labeled for two PNN markers, the PNN CSPG aggrecan (anti-ACAN) and the PNN CS-GAGs (WFL), and co-localized with a neuronal marker (anti-NeuN) marker (**Table II**). First, sections were rinsed thrice in PBS and then incubated overnight at 4°C under constant agitation with the appropriate antibody or lectin diluted in a blocking solution of PBS containing 0.2% Triton-X and 2% normal goat serum. Mouse tissues had an additional 1 h of blocking with 10% normal goat serum in PBS with 0.2% Triton-X before primary labeling. After overnight primary incubation, sections were rinsed thrice in PBS and incubated for 2h at RT with the appropriate fluorophore-conjugated secondary antibody or streptavidin (**Table II**) diluted in the blocking solution. Next, tissues were rinsed and incubated with TrueBlack® (Biotium, #23007, Fremont, California, USA) for 80 seconds to remove endogenous autofluorescence from lipofuscin and other cellular debris. Sections were then mounted on Superfrost charged slides and coverslipped with Vectashield mounting medium with DAPI (Vector Laboratories, H-1800, Newark, California, USA) and stored at 4°C until imaged on an Evident Scientific VS120 Slide Scanner at 20X magnification.

**Table II:** Specific labeling details.

| **Table II: Specific labeling details** |  |  |  |  |  |
| --- | --- | --- | --- | --- | --- |
| **Abbreviation** | **Primary** | **Labels** | **Target species** | **Product** | **Fluorophore conjugated-secondary** |
| *Human samples* |  |  |  |  |  |
| **WFL/WFA** | Biotinylated Wisteria Floribunda Lectin | Perineuronal nets | Human | Vector Laboratories, B-1355, 1:2500 | Alexa 488-Streptavidin, Jackson ImmunoResearch, 016-540-084, 1:500 |
| **NeuN** | Anti-neuronal nuclei | Neurons | Human | Abcam, ab104225, 1:1000 | DyLight-594 anti-rabbit, Jackson ImmunoResearch, 111-515-144, 1:500 |
| **ACAN** | Anti-chondroitin sulfate proteoglycan brain core protein (Cat-301) | Perineuronal nets | Human | Millipore, MAB5284, 1:500 | Alexa-647 anti-mouse, Jackson ImmunoResearch, 115-605-146, 1:500 |
| *Mouse samples* |  |  |  |  |  |
| **WFL/WFA** | Biotinylated Wisteria Floribunda Agglutinin | Perineuronal nets | Mouse | Vector Laboratories, B-1355, 1:1500 | Cy3-Streptavidin, Jackson ImmunoResearch, 016-160-084, 1:500 |
| **NeuN** | Anti-neuronal nuclei | Neurons | Mouse | Millipore, MAB377, 1:500 | Alexa-647 anti-mouse, Jackson ImmunoResearch, 115-605-146, 1:500 |
| **ACAN** | Anti-aggreccan | Perineuronal nets | Mouse | Millipore, AB1031, 1:100 | Alexa-488 anti-rabbit. Jackson ImmunoResearch, 111-485-144, 1:500 |

Image analysis was conducted with QuPath (Bankhead, P., Loughrey, M.B., et al. 2017) version 0.3.2. Two to four ROIs spanning all cortical or subcortical layers were counted per human sample in the vmPFC (CTRL = 11, DS = 9, DS-CA = 9), EC (CTRL = 8, DS = 16, DS-CA = 14), HPC (CTRL = 8, DS = 15, DS-CA = 16) or mouse sample in the mPFC, EC, vHPC (CTRL = 10, ELS = 9). ROIs were manually investigated in 20x images by a blinded researcher who manually classified PNNs as double positive (WFL+/ ACAN+) or solely labeled by WFL+ or ACAN+. Note, aggrecan labeling was only classified as a PNN when it was localized surrounding the cell body of a neuron, i.e. if it was not 100% overlapping with intracellular neuronal staining (NeuN). A total of 30982 PNNs were manually classified in the human brain (14,644 in vmPFC, 6,920 in EC, 9,418 in HPC) and 5,902 in the mouse brain (1,419 in mPFC, 2,233 in EC, 2,250 vHPC). PNN counts were normalized to 100 allowing for comparison between groups using percentage of total PNNs classified per brain region.

### Statistical analyses

Statistical tests were conducted using Prism version 10 (GraphPad Software, Boston, Massachusetts, USA) and SPSS version 29.0 (IBM Corp, Armonk, New York, USA). Distribution and homogeneity of variances were assessed through Shapiro-Wilk and Levene’s tests, respectively. Spearman’s correlation was used to explore the relationship between dependent variables and covariates (age, PMI, pH, types of antidepressants and number of medications) (**Supplementary Figure 1**). Age exhibited a correlation with PNN labeling pattern, prompting the use of analysis of covariance (ANCOVA) to compare human labeling patterns. Analysis of variance (ANOVA) was used to compare sulfation pattern. Statistical tests were all two-sided with a significance threshold of 0.05. Data are presented as mean ± standard error of the mean unless otherwise indicated.

## Acknowledgments

We extend our heartfelt appreciation to the families of the donors who graciously agreed to donate the brains of their beloved family members. The present study used the services of the Molecular and Cellular Microscopy Platform at the Douglas Hospital Research Centre. Mass spectrometry work was supported by University of Washington School of Pharmacy’s Mass Spectrometry Center.

## Abbreviations

ACAN: aggrecan
ANCOVA: analysis of covariance
ANOVA: analysis of variance
C4ST-1: chondroitin 4-sulfotransferase-1
C6ST-1: chondroitin 6-sulfotransferase-1
CCAC: Canadian Council on Animal Care
CECA: Childhood Experience of Care and Abuse
chABC: chondroitinaseABC
CS-GAG: chondroitin sulfate glycosaminoglycan
CS: chondroitin sulfate
CSPGs: chondroitin sulfate proteoglycans
CTRL: healthy individuals having died suddenly
DBCBB: Douglas-Bell Canada Brain Bank
DS-CA: depressed suicides with a history of child abuse
DS: depressed suicides without a history of child abuse
DSM-IV: Diagnostic and Statistical Manual of Mental Disorders
EC: entorhinal cortex
ECM: extracellular matrix
ELS: early-life stress
ESI: electrospray ion
GalNAc: N-acetylgalactosamine
GlcA: D-glucuronic acid
HAS: hyaluronan synthase
HPC: hippocampus
LC-MS/MS: liquid chromatography tandem mass spectrometry
mPFC: medial prefrontal cortex
Otx2: orthodenticle homeobox protein 2
P: postnatal day
PBS: phosphate buffered saline
PFA: paraformaldehyde
PNNs: perineuronal nets
PV: parvalbumin
ROI: regions of interest
RT: room temperature
TQ: triple quadrupole
UPLC: ultra-performance liquid chromatographic system
vHPC: ventral hippocampus
vmPFC: ventromedial prefrontal cortex
WFL: Wisteria Floribunda Lectin

## Data Availability

All data generated for this study are contained within the manuscript. For further queries, the corresponding author NM may be contacted.

## Author Contributions

CB, KA and NM conceived the study. GT participated in the acquisition and clinical characterization of the brain samples. CB, ST, SN, CH and MAD contributed to immunohistological experiments. RR, GF and BG conducted animal work. KA, AH and KH conducted LC-MS/MS experiments. CB conducted data analysis. CB, NM, KA prepared the manuscript and all authors contributed to and approved its final version.

