## Supplementary Figure 1 for "Chondroitin sulfate glycan sulfation patterns influence histochemical labeling of perineuronal nets: a comparative study of interregional distribution in human and mouse brain"

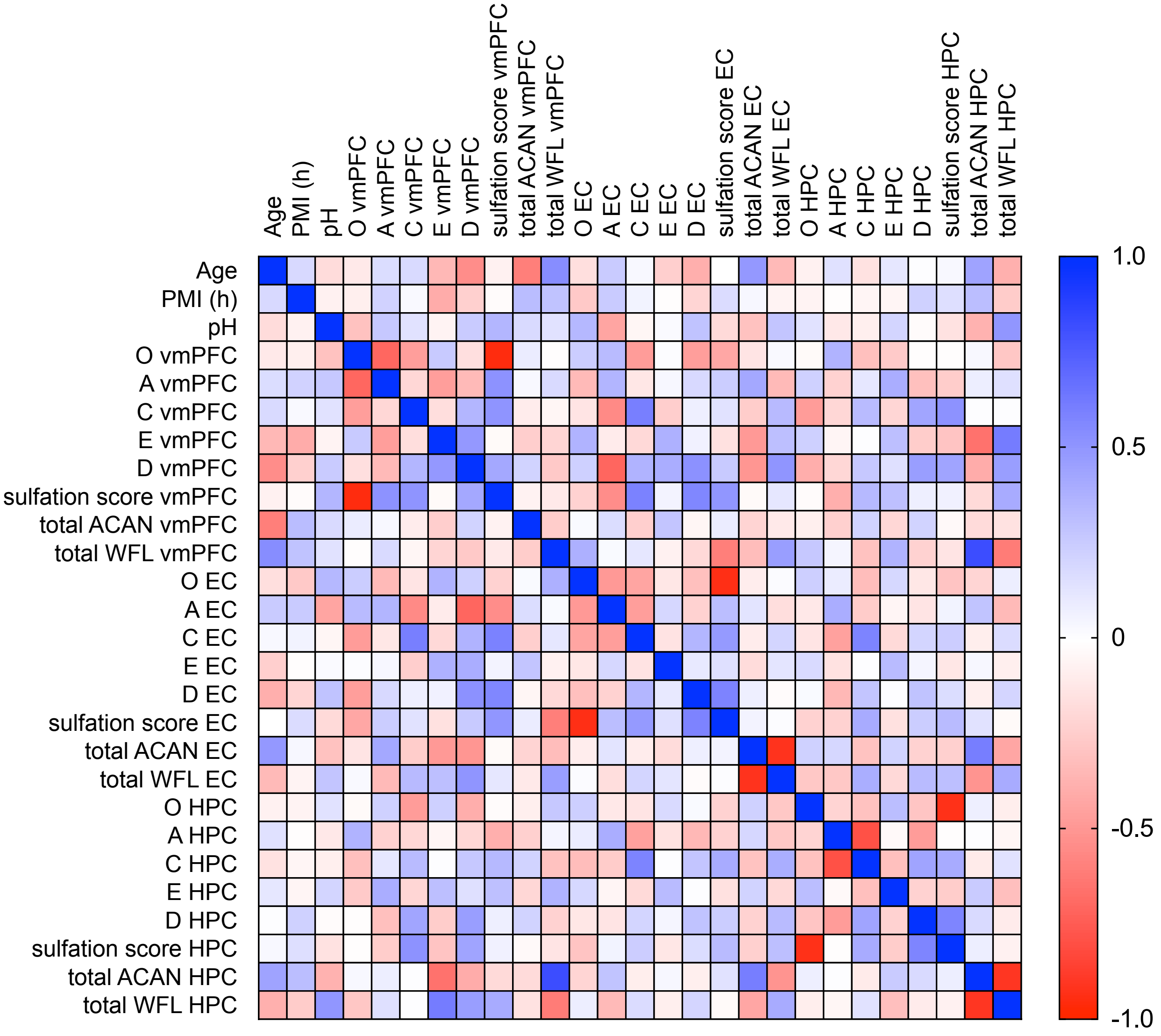


**Supplementary Figure 1**: Correlation matrix demonstrating Spearman's correlation between dependent variables and covariates (age, PMI and pH) in human samples. Each cell in the matrix contains the correlation coefficient corresponding to the pairwise comparison of variables. Correlation coefficients range from -1 to +1, where values closer to +1 indicate a strong positive correlation (blue), values closer to -1 indicate a strong negative correlation (red) and values around 0 indicate no correlation. The color intensity or shading of each cell indicates the strength and direction of the correlation, with darker shades representing stronger correlations.
