## Supplementary Table for "Chondroitin sulfate glycan sulfation patterns influence histochemical labeling of perineuronal nets: a comparative study of interregional distribution in human and mouse brain"

| Table 1: Human sulfation controls | | | | |
|---|---|---|---|---|
| Isomer | | vmPFC | EC | HPC |
| CS-A | Mean % | 77.6 | 76.2 | 81.2 |
| | SD | 1.9 | 1.6 | 1.2 |
| CS-O | Mean % | 13.6 | 11.4 | 5.0 |
| | SD | 2.2 | 0.6 | 0.9 |
| CS-D | Mean % | 2.0 | 2.8 | 3.1 |
| | SD | 0.5 | 0.6 | 0.4 |
| CS-C | Mean % | 5.6 | 8.4 | 9.5 |
| | SD | 1.7 | 1.4 | 1.1 |
| CS-E | Mean % | 1.2 | 1.1 | 1.1 |
| | SD | 0.4 | 0.2 | 0.2 |

| Table 2: Mouse sulfation controls | | | | |
|---|---|---|---|---|
| Isomer | | mPFC | EC | vHPC |
| CS-A | Mean % | 70.0 | 73.9 | 79.6 |
| | SD | 1.9 | 1.1 | 3.2 |
| CS-O | Mean % | 17.6 | 16.8 | 8.0 |
| | SD | 1.5 | 0.7 | 1.4 |
| CS-D | Mean % | 2.2 | 1.4 | 1.7 |
| | SD | 0.3 | 0.2 | 0.4 |
| CS-C | Mean % | 8.3 | 6.4 | 9.2 |
| | SD | 0.8 | 0.6 | 1.4 |
| CS-E | Mean % | 1.4 | 0.9 | 0.7 |
| | SD | 0.08 | 0.1 | 0.1 |

| Table 3: Human labeling controls | | | | | | | | |
|---|---|---|---|---|---|---|---|---|
| | | vmPFC | EC | HPC | | vmPFC – EC | vmPFC – HPC | EC - HPC |
| ACAN | Mean % | 23.4 | 20.5 | 54.2 | p-value | 1.0 | <0.001 | < 0.001 |
| | SD | 8.7 | 15.8 | 15.8 | | | | |
| WFL | Mean % | 28.3 | 66.8 | 25.7 | p-value | < 0.001 | 1.0 | < 0.001 |
| | SD | 10.1 | 19.4 | 13.7 | | | | |
| both | Mean % | 48.3 | 12.7 | 20.1 | p-value | < 0.001 | < 0.001 | 0.3 |
| | SD | 8.8 | 10.2 | 6.9 | | | | |

| Table 4: Mouse labeling controls | | | | | | | | |
|---|---|---|---|---|---|---|---|---|
| | | mPFC | EC | vHPC | | mPFC – EC | mPFC – vHPC | EC - vHPC |
| ACAN | Mean % | 0 | 0.6 | 12.5 | p-value | 0.3 | 0.0002 | 0.0002 |
| | SD | 0 | 0.7 | 4.1 | | | | |
| WFL | Mean % | 94.7 | 92.3 | 39.5 | p-value | > 0.9999 | < 0.0001 | < 0.0001 |
| | SD | 5.8 | 4.9 | 9.2 | | | | |
| both | Mean % | 5.5 | 7.1 | 47.9 | p-value | > 0.9999 | < 0.0001 | < 0.0001 |
| | SD | 5.8 | 4.6 | 10.2 | | | | |

Mean %: Mean percentage of PNNs with this labeling
SD: Standard deviation
